# An unbiased, automated platform for scoring dopaminergic neurodegeneration in *C. elegans*

**DOI:** 10.1101/2023.02.02.526781

**Authors:** Andrew S. Clark, Zachary Kalmanson, Katherine Morton, Jessica Hartman, Joel Meyer, Adriana San-Miguel

**Affiliations:** Chemical and Biomolecular Engineering, North Carolina State University, Raleigh, North Carolina, USA; Nicholas School of the Environment, Duke University, Durham, North Carolina, USA; Biochemistry and Molecular Biology, Medical University of South Carolina, Charleston, South Carolina, USA

## Abstract

*Caenorhabditis elegans* (*C. elegans*) has served as a simple model organism to study dopaminergic neurodegeneration, as it enables quantitative analysis of cellular and sub-cellular morphologies in live animals. These isogenic nematodes have a rapid life cycle and transparent body, making high-throughput imaging and evaluation of fluorescently tagged neurons possible. However, the current state-of-the-art method for quantifying dopaminergic degeneration requires researchers to manually examine images and score dendrites into groups of varying levels of neurodegeneration severity, which is time consuming, subject to bias, and limited in data sensitivity. We aim to overcome the pitfalls of manual neuron scoring by developing an automated, unbiased image processing algorithm to quantify dopaminergic neurodegeneration in *C. elegans*. The algorithm can be used on images acquired with different microscopy setups and only requires two inputs: a maximum projection image of the four cephalic neurons in the *C. elegans* head and the pixel size of the user’s camera. We validate the platform by detecting and quantifying neurodegeneration in nematodes exposed to rotenone, cold shock, and 6-hydroxydopamine using 63x epifluorescence, 63x confocal, and 40x epifluorescence microscopy, respectively. Analysis of tubby mutant worms with altered fat storage showed that, contrary to our hypothesis, increased adiposity did not sensitize to stressor-induced neurodegeneration. We further verify the accuracy of the algorithm by comparing code-generated, categorical degeneration results with manually scored dendrites of the same experiments. The platform, which detects 19 different metrics of neurodegeneration, can provide comparative insight into how each exposure affects dopaminergic neurodegeneration patterns.

## I. Introduction

Neurodegenerative disorders like Parkinson’s disease (PD) are an increasingly significant health crisis in part due to aging populations. PD is characterized by the progressive degeneration of dopaminergic neurons in the substantia nigra region of the brain(1). Decades of research have examined factors contributing to its pathology, identifying causative roles for several genetic mutations and environmental toxicants(2). PD takes two forms : familial and idiopathic. Familial PD provides insight into the mechanisms of this disease, with known mutations in genes including LRRK2, PINK1 and PRKN all increasing disease likelihood(3). However, up to 90% of cases are idiopathic(4). PD currently has no cure, underscoring the importance of identifying the root causes of PD and identifying potential targets for pharmaceutical intervention.

The risk of developing idiopathic PD has been correlated with exposure to environmental toxicants such as the pesticides paraquat and rotenone(5). Unfortunately, there are tens of thousands of high-volume synthetic chemicals in use, and very few have been tested for their potential to cause neurodegeneration(6,7). Furthermore, with known genetic vulnerabilities increasing PD risk, and many unknowns regarding how these genetic and environmental factors interact, the ability to screen chemicals alone is not enough. With the multitude of gene-environment interactions to consider, high-throughput screening will be essential for evaluating chemical potential to induce dopaminergic neurodegeneration. Currently, there is not a widely accepted high-throughput screening method to address this concern. The slow growth, expensive maintenance, and extensive work required to investigate neurodegeneration in mammalian models has led to the exploration of models that are more adaptable to higher-throughput methods. Cell culture models such as the commonly used SHSY-5Y cells have been employed but cannot account for the complex intracellular and intercellular relationships that occur *in vivo*. In addition, these neurons do not form their natural networks and the role of their dysfunction cannot be contextualized to an organismal scale(8). Here, we propose the use of a small transparent nematode, *Caenorhabditis elegans* (*C. elegans*) and a novel automated system for quantifying dopaminergic neurodegeneration to increase the throughput, decrease the bias, and improve the accuracy of the data collected in *C. elegans* dopaminergic neurodegeneration experiments.

*C. elegans* is a well-studied model for examining factors that can lead to dopaminergic neurodegeneration(9–14). This small nematode with a rapid life cycle and short life span makes examination of aging and aging-related pathologies such as neurodegeneration simple, and its transparent body makes fluorescence imaging in living animals feasible. *C. elegans* possesses a fully characterized nervous system with eight dopaminergic neurons, of which the four cephalic (CEP) neurons have a particularly well-studied pattern of damage characterized by specific degenerative phenotypes including blebbing and breakage(9,15). *C. elegans* demonstrate well conserved aging and metabolic pathways, both critical to the pathology of PD, and the CEP neurons have replicated degenerative effects induced by common PD models, such as rotenone and 6-OHDA treatment(13,15,16).

Many different systems for analyzing or scoring degeneration of the CEP neurons have been proposed. Some look at one marker of degradation, such as loss of the neuronal cell body(17) or quantification of the loss of fluorescence in the soma(18,19). Though easily quantified, this metric may be influenced by the genetic or chemical influences on the promoter used to drive fluorescent protein expression. Other methods attempt to quantify damage within the dendrites of the CEP neurons, with varying degrees of specificity. Typically, this is completed by categorizing either using a binary approach (damaged or healthy) or a more granular grading system, such as 5-point or 7-point systems that grade the dendrites based on a semi-quantitative analysis of morphological features(14,15,20,21). These scoring systems provide more nuance and can help assess changes in neuron health across different doses of toxins, but they are limited by the subjective, qualitative nature of manual scoring. A purely quantitative analysis of neuron images could better reveal subtle differences in degradation(22). For example, it may allow for investigation of if and how age-related deterioration may present differently than deterioration due to toxicants, or whether degradation from exposure to one class of toxin may present different morphological effects than exposure to another. Thus, it is of significant interest to develop a platform capable of portraying a comprehensive and precise assay of the neuron’s health.

Manual scoring of neurons has other drawbacks, such as bias. While there are techniques and software that can be used to ensure consistency across scoring groups(23), the possibility of bias is inherent to manual observation. Another issue is the time and labor that comes with scoring large sets of images by hand. In addition, it can be difficult for the human eye to consistently pick up on subtle morphological irregularities, defined here as features. For instance, information like the size or shape of a feature of interest is not measurable by eye yet may yield important information about the neuron’s health.

Computer vision algorithms for tracking and analyzing neurons have attempted to alleviate many of these issues(24,25). However, it can be difficult for an algorithm to correctly identify dendrites due to breaking segments and overlaps. Many projects turn to machine learning models, specifically Convolutional Neural Networks, to aid this(26,27). Training machine learning models requires large quantities of data at the risk of overfitting the model to the training set. For example, a model trained on images from one microscope or image setting may struggle when given data from a different set up. Building a platform robust enough to handle data from multiple sources would help standardize further research using the *C. elegans* model.

We present an Automated, Unbiased Dopaminergic Degeneration Identification Tool (AUDDIT), a new algorithmic approach to trace and score CEP dendrites without the need for user training. Our approach can segment images to identify dendrites, track and analyze the dendrites to assess their health, and pick up morphological features such as blebbing. AUDDIT can analyze images from different imaging setups, which may vary in camera resolution, intensity, and pixel size. Compatibility across differing imaging setups can help lower the barriers of entry into image processing based quantitative neurodegenerative analysis. We verify the algorithm’s versatility by quantifying dopaminergic neurodegeneration in nematodes exposed to rotenone, cold shock, and 6-hydroxydopamine (6-OHDA) using 63x epifluorescence, 63x confocal, and 40x epifluorescence microscopy, respectively, while also using different light sources and cameras for acquisition. Through the quantitative data produced by this algorithm, further research can be undertaken to highlight subtle, yet distinct, dopaminergic degradation phenotypes in a consistent and comprehensive manner.

## II. Materials and Methods

### A. Strains used

For imaging CEP neurons in the head region, the BY200 [vtIs1 (*dat*-1p::GFP, *rol*-6)] strain was utilized. The 6-OHDA exposures used RB1600 [*tub-1*(ok192)[, and RB2237 [*tub-2*(ok3024)[, which were crossed with BY200. Worms were cultured on K-agar(28) plates seeded with OP50 *E. coli* strain OP50 at 20 °C. Additional culturing details specific to different exposure scenarios are provided below.

### B. Rotenone exposure

Worm populations were age-synchronized by extracting worm eggs from gravid adult worms using a 1% bleach and 0.1 M NaOH solution(29). Worms were then grown from the L1 stage in 1 mL wells filled with complete K medium (2.36 KCl, 3 g NaCl, 5 mg cholesterol, 1 mL 1M CaCl_2_, 1 mL 1M MgSO_4_ in 1 L of water)(28). Worms were kept well fed using bacterial strain HB101 at 50 mg/ml. Well plates were gently rocked and incubated at 25 °C. Rotenone diluted in DMSO was dosed appropriately at either 0 µM, 0.03 µM, or 0.5 µM and added to the wells at hatching. Rotenone concentrations were re-dosed daily to account for absorption and chemical breakdown. Dilutions were made ensuring final DMSO concentration in the K medium was 0.125% by volume. Worms were removed from the wells once reaching the L4 stage (an extra 24 h was required for worms to reach L4 in 0.5 µM rotenone), then transferred to slides for prompt imaging (see microscopy and imaging) .

### C. Cold shock exposure

Animals were age-synchronized by extracting worm eggs from gravid adult worms using a 1% bleach and 0.1 M NaOH solution(29). To improve age-synchronization, extracted eggs were then placed in M9 buffer (3 g KH_2_PO_4_, 6 g Na_2_HPO_4_, 5 g NaCl, and 1 mL of 1 M MgSO_4_ in 1 L of water) overnight. Newly hatched L1 worms were then transferred to a solid nematode growth media (NGM)(30) plate seeded with *Escherichia coli* OP50. Worm populations were then cultured at 25 °C until they reached day 1 of adulthood. A subpopulation of worms was then imaged (see microscopy and imaging) prior to the 16-hour cold shock. Worms were then transferred to the 4 °C refrigerator for 16 hours. Following the cold shock, plates were placed at room temperature for 1 hour prior to imaging.

### D. 6-hydroxydopamine exposures

Worms were age-synchronized by transferring adults to K-agar plates seeded with OP50 for 3 hours and allowed to lay eggs. Afterwards, all worms were washed from the plates with K medium. The remaining eggs were allowed to hatch and age to adulthood. Beginning on day 1 of adulthood, they were transferred to fresh plates each day to separate them from progeny until adult day 5. On adult day 5 worms were exposed in K+ medium to either a vehicle control of ascorbic acid, or 6-hydroxydopamine (6-OHDA) in ascorbic acid for one hour. After exposure, animals were washed three times with 2 mL K-medium with 2 minutes for gravity settling between addition of K-medium and aspiration. They were subsequently plated on K-agar plates seeded with OP50 where they remained for 48 hours until imaging.

### E. Microscopy and imaging

Animals were placed on a glass slide with 2% agarose pads. To immobilize worms for high-resolution imaging, two methods were used. For the 6-OHDA exposure, worms were picked via stainless steel pick onto 2% agarose pads on glass slides and paralyzed with 500 mM sodium azide. Alternatively, for cold shock imaging, worms were placed in a solution of 5 mM tetramisole and M9 buffer prior to transferring to the agarose pad on a glass slide. Worms naturally move in a lateral orientation that causes the four CEP neurons to overlap while imaging. Thus, immobilizing prior to mounting worms increased worm orientation variability and improved the chances that all CEP neurons in the head would be visible in a maximum projection image. The experimental exposure determined how the images were acquired. For the rotenone exposure experiments, images were acquired with an inverted Leica DMi8 widefield fluorescence microscope, a Hamamatsu Orca-Fusion camera using a 63x objective (NA = 1.40), and a metal halide light source. Worms from the cold shock experiments were imaged with an inverted Leica DMi8 widefield fluorescence microscope equipped with a spinning disk confocal head (Crest Optics X-light V2), a Hamamatsu Orca-Fusion camera using a 63x objective (NA = 1.40), and a Laser Diode Illuminator (89 North LDI). Z-stack images were acquired using a 0.75 µm step size and 35 steps. Lastly, worms from the 6-OHDA experiments were imaged with a Keyence BZ-X710 fluorescence microscope using a 40x objective. Z-stacks images were acquired using a 0.5 µM step size.

### F. System requirements and performance

AUDDIT v1.0 was scripted in MATLAB 2021b (MathWorks) with the Image Processing and Statistics toolboxes installed. The graphical user interface (GUI) was developed using the built-in App Designer. The GUI works as a standalone desktop application and does not require a MATLAB license. Both the GUI and the MATLAB scripts can be found on GitHub: https://github.com/zkalmanson/AUDDIT. For the data in this article, AUDDIT v1.0 was used. We plan to continue to improve performance of AUDDIT and will upload new updates to the GitHub link as they occur.

The software package was tested on a desktop computer with the Apple Silicon M1 chip and 16 GB memory. On 16-bit images captured with the Hamamatsu Orca-Fusion (2304 x 2304 pixels), AUDDIT could analyze images at ∼10 seconds per image. On the smaller, 8-bit images captured on the Keyence BZ-X700 (760 x 920 pixels), it could analyze images at ∼2 seconds per image. It is important to note that displaying images while the algorithm is running significantly increases the processing time.

### G. Statistical analysis

The statistical analysis throughout this study used the Statistics and Machine Learning Toolbox in MATLAB 2021b. For each exposure, each dendrite analyzed is counted as N = 1. Each exposure and metric were analyzed with a one-way ANOVA. For individual exposure analysis (Figs 3-5), the Bonferroni multiple comparison correction was used(31). When comparing exposure patterns to the respective controls in Fig 6, the Dunnett’s correction was used(32). All statistical output with related p values for each experiment are available in the Supplementary, as well as the Dryad repository.

## III. Results

### A. Unbiased, automatic algorithm for neurodegeneration detection

AUDDIT aims to increase throughput and decrease bias in obtaining quantifiable neurodegeneration metrics. We sought to limit user input to a maximum projection image of the four fluorescently tagged CEP neurons in the *C. elegans* head and the pixel size of the user’s camera. The pixel size of the user’s camera is used throughout the algorithm to scale functions that are size-dependent, such as creating structuring elements or determining minimum separation between points. Once the pixel size is set and images are loaded, the algorithm completes a series of steps to detect, track, and report neurodegeneration metrics (Fig 1).

**Fig 1.**
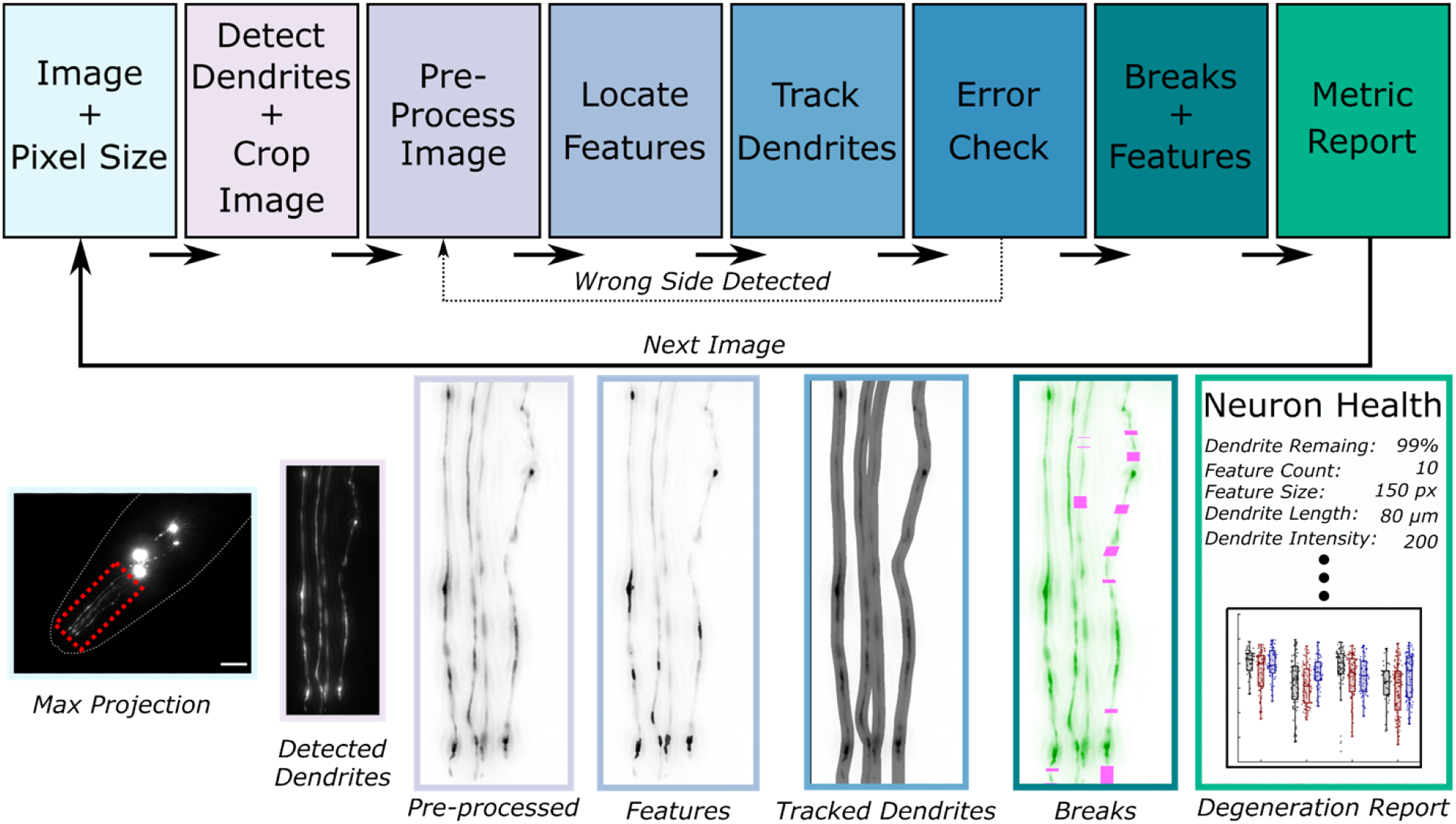
Overall neurodegeneration algorithm schematic. The algorithm requires the user to provide maximum intensity projection image(s) of the CEP neurons and the pixel size in microns. Once the images and pixel size are set, the algorithm automatically detects dendrite locations and crops the image accordingly. Next, the cropped image undergoes a preprocessing step, which permits feature detection and dendrite tracking. An error check follows this step to ensure the dendrites were correctly identified. Using the tracks and feature locations, dendrite breakage and feature metrics are quantified and stored for an overall neurodegeneration report. Max projection scale bar = 15 µm.

### i. Dendrite detection

Because they show degeneration earlier than the cell bodies, dendrites are often the focus when studying neurodegeneration in CEP neurons (15,20,33–35). Thus, the first step of the algorithm automatically crops the images such that only the dendrites are in view. This is feasible because the CEP dendrites, when correctly oriented, present a common morphology of four long, vertical strands. Preprocessing uses an initial global contrast adjustment to ensure the dendrites are bright enough to be detected, as their fluorescence is significantly lower than the cell body of the neuron. Since the cell bodies are often the highest intensity pixels, a binarization step with a threshold level aimed to only detect the brightest pixels is performed (Fig 2A). Once the cell body is detected, the image is rotated such that the dendrites are aligned with the vertical axis. The image is then split horizontally into two smaller images using the bounding box of the cell body as the reference point for cropping (Fig 2A). To determine which of the newly created images contains the dendrites, another binarization step with a less stringent threshold level is completed on both images. The resulting binarized images contain regions of dendrites and any other objects in the frame (Fig 2A). Various properties of the identified objects, such as orientation, major axis length, circularity, and area, are extracted using the built-in regionprops() function in MATLAB. Next, to determine which image contains the dendrites, the region properties from the split images are compared. For example, the cell bodies tend to be circular, whereas the dendrites tend to be elongated and oriented along the vertical axis. Thus, the image with fewer circular objects is chosen as the image containing dendrites. Finally, once the correct side of the image is determined, border removal functions are used to exclude the cell bodies and any unwanted objects around the edges of the image. The image is further cropped to remove the extra space vacated by unwanted object removal (Fig 2A).

**Fig 2.**
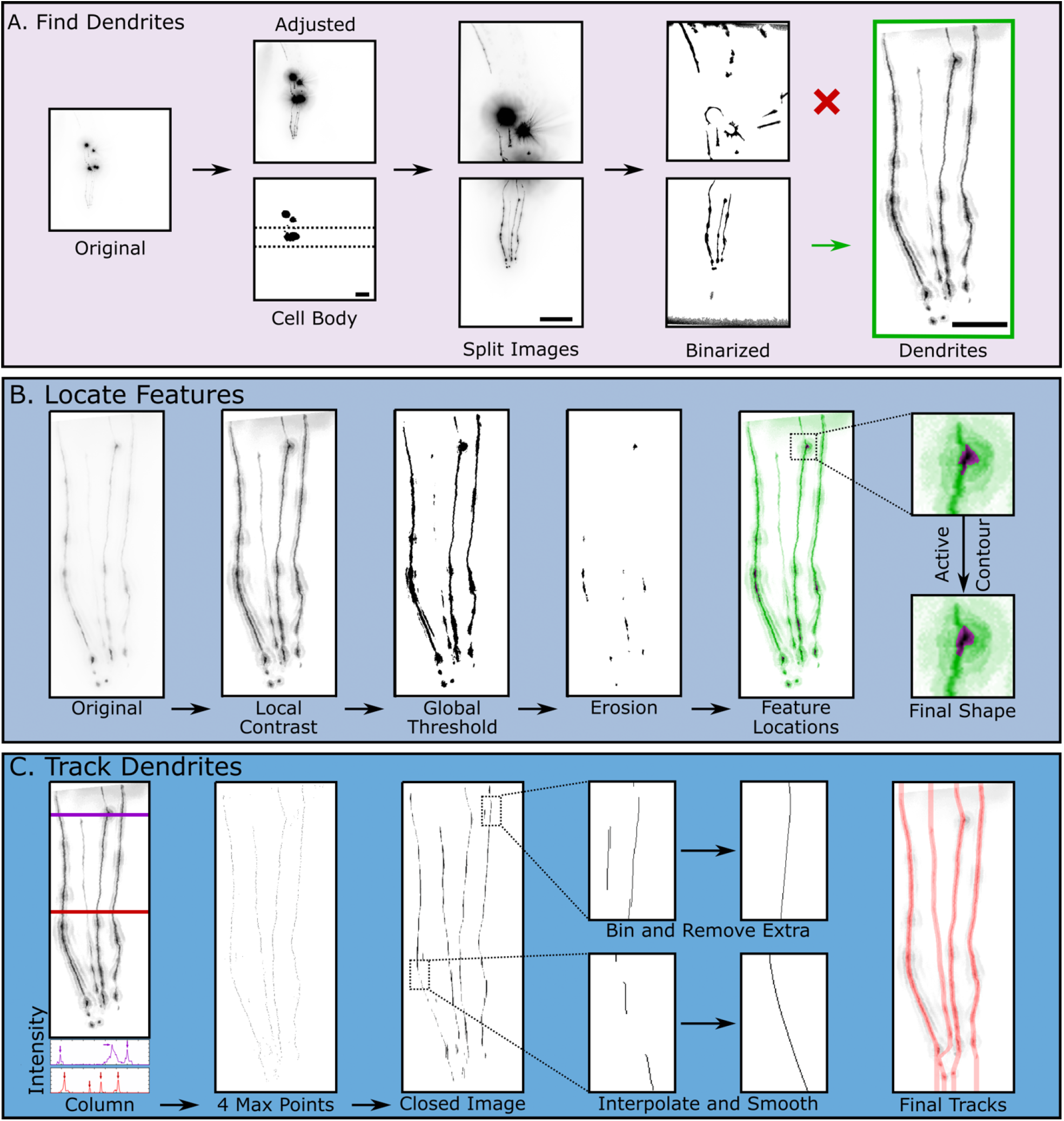
Key steps in neurodegeneration detection algorithm. **A**. Dendrite detection steps. The original image is first contrast adjusted and binarized to detect the location of the cell body (Scale bar = 25 µm). Next, the image is split into two images (Scale bar = 25 µm). Using the structures obtained through binarizing these images, the algorithm detects which image contains the dendrites (Scale bar = 10 µm). **B**. Feature detection steps. First, the cropped image undergoes local contrast adjustment to improve the subsequent global thresholding and binarization step. Feature locations are obtained using horizontal image erosion. Lastly, active contour segmentation of these features improves feature shape definition. **C**. Dendrite tracking. The local contrast adjusted image is used as a starting point to track dendrites. Each row of pixel intensities is analyzed to obtain 4 local maximum points, which correspond to the four dendrites. The resulting binary image is then closed with a vertical structuring element to connect points where no local maxima is detected. Next, for each row, every foreground point is binned and labeled into its respective dendrite and extra foreground points are removed. Lastly, rows with missing dendrites are interpolated and smoothed to create the final tracks

**Fig 3.**
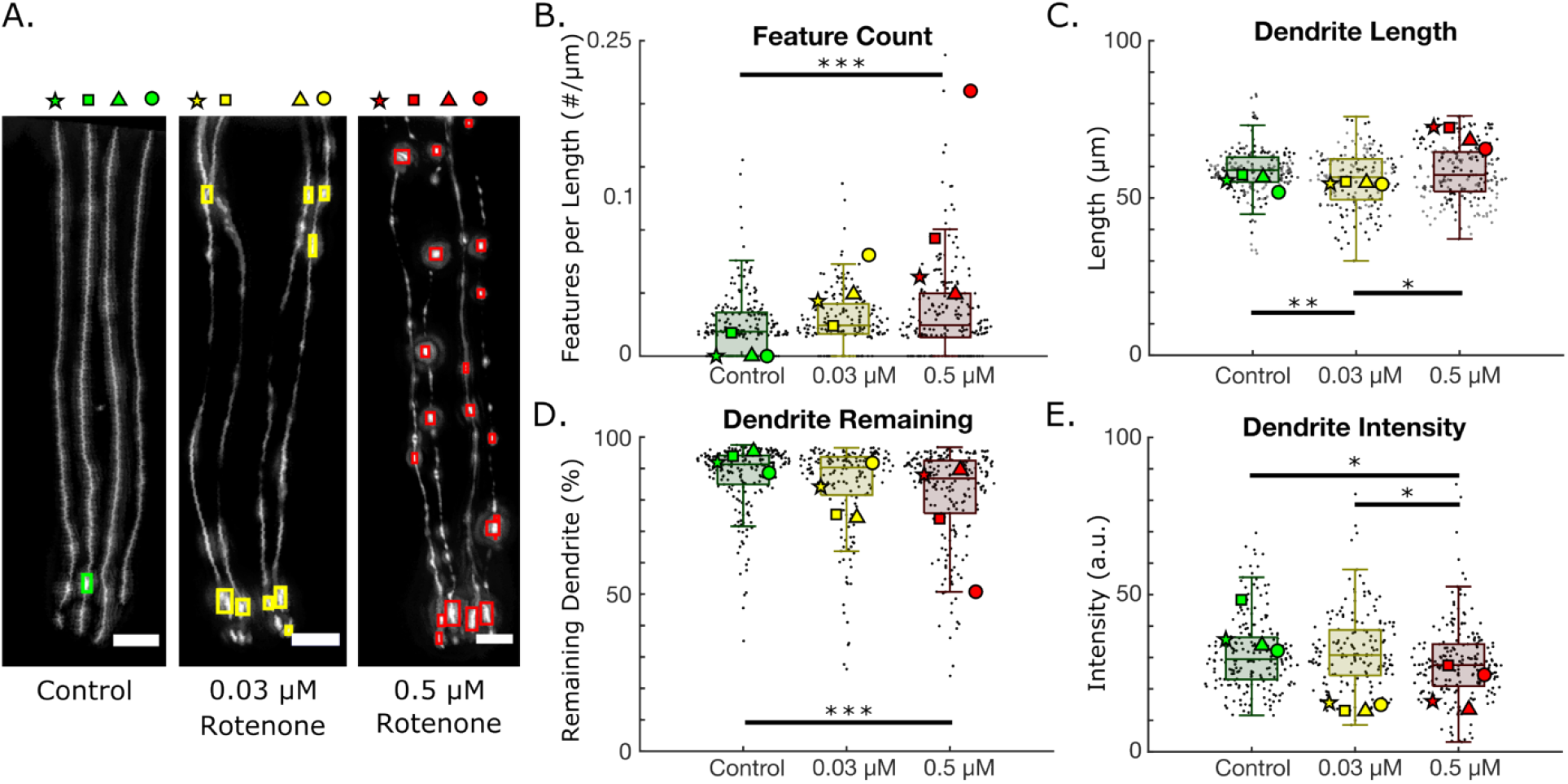
Neurodegeneration induced by rotenone exposure in BY200 worms. A. Example images and detection of (i) control, (ii) 0.03 µm rotenone exposed, and (iii) 0.5 µM rotenone exposed worms (scale bar = 10 µm). Each dendrite is labeled as a star, square, triangle, or circle. N=1 represents a single dendrite. B-E. Selected neurodegeneration metrics for control (N = 204), 0.03 µM rotenone (N = 168), and 0.5 µM rotenone exposed worms (N = 208). B. Number of detected features (blebs) per dendrite length. (0.03 µM rotenone to 0.05 µM rotenone; p = 0.066). C. Dendrite lengths obtained from tracked dendrites D. Percentage of dendrite remaining and E. Dendrite intensities. The neurodegeneration results for each dendrite in panel (A) are overlaid. (One-way ANOVA. *p < 0.05; **p < 0.01; ***p < 0.001).

**Fig 4.**
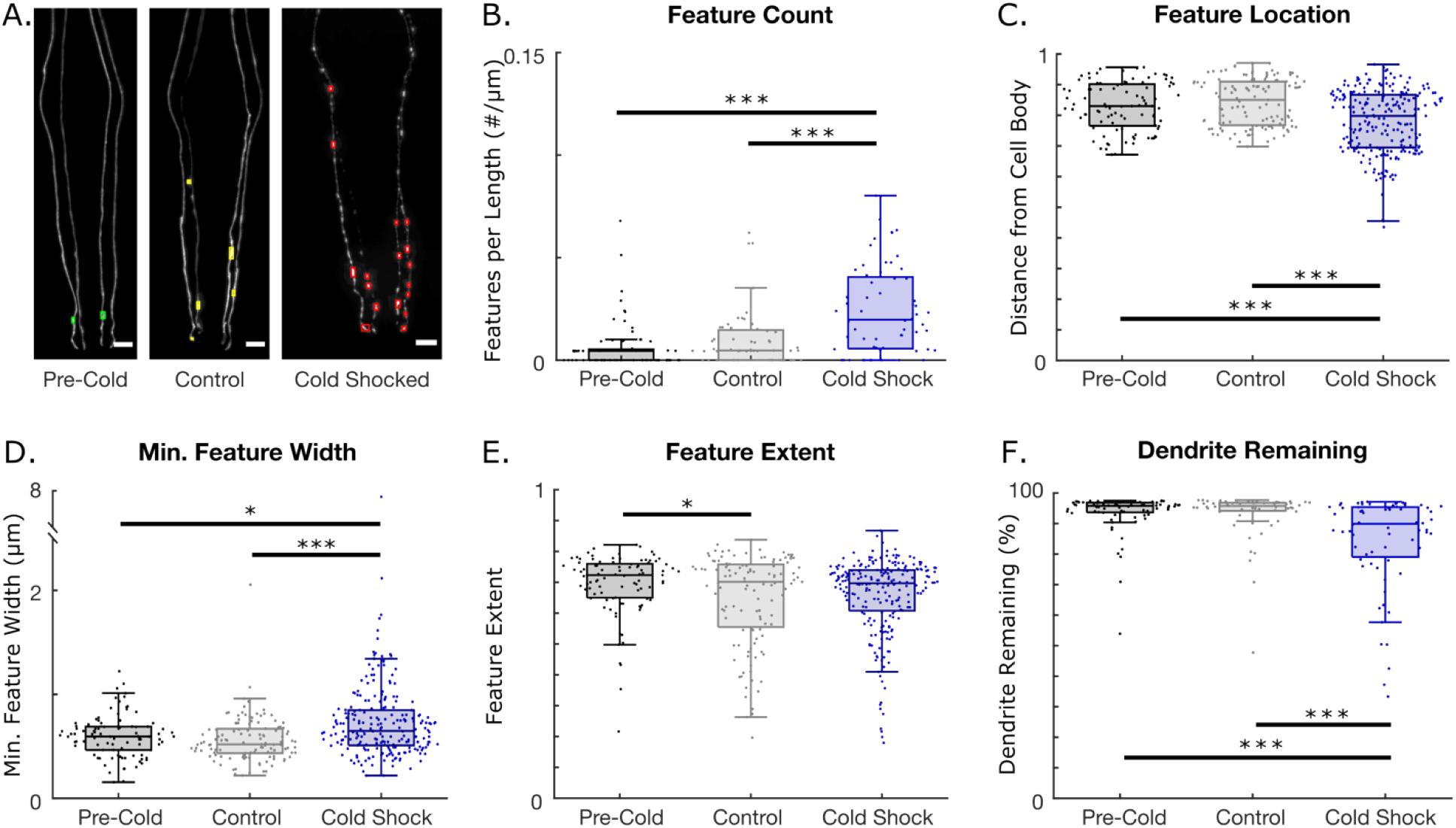
Neurodegeneration induced by 16-hour cold shock at 4°C. N=1 represents a single dendrite. A. Example confocal images of CEP neurons and detected features for pre-cold shock, control, and cold shocked worms (Scale bar = 10 µm). B-F. Neurodegeneration metrics for pre-cold shock (N = 72), control (N = 56) and cold-shocked worms (N = 68). B. Feature count per dendrite length. C. Normalized feature location and distance from the cell body. D. Minimum caliper diameters of detected features. E. Feature extent. F. Percentage of dendrite remaining. (One-way ANOVA. *p < 0.05; **p < 0.01; ***p < 0.001).

**Fig 5.**
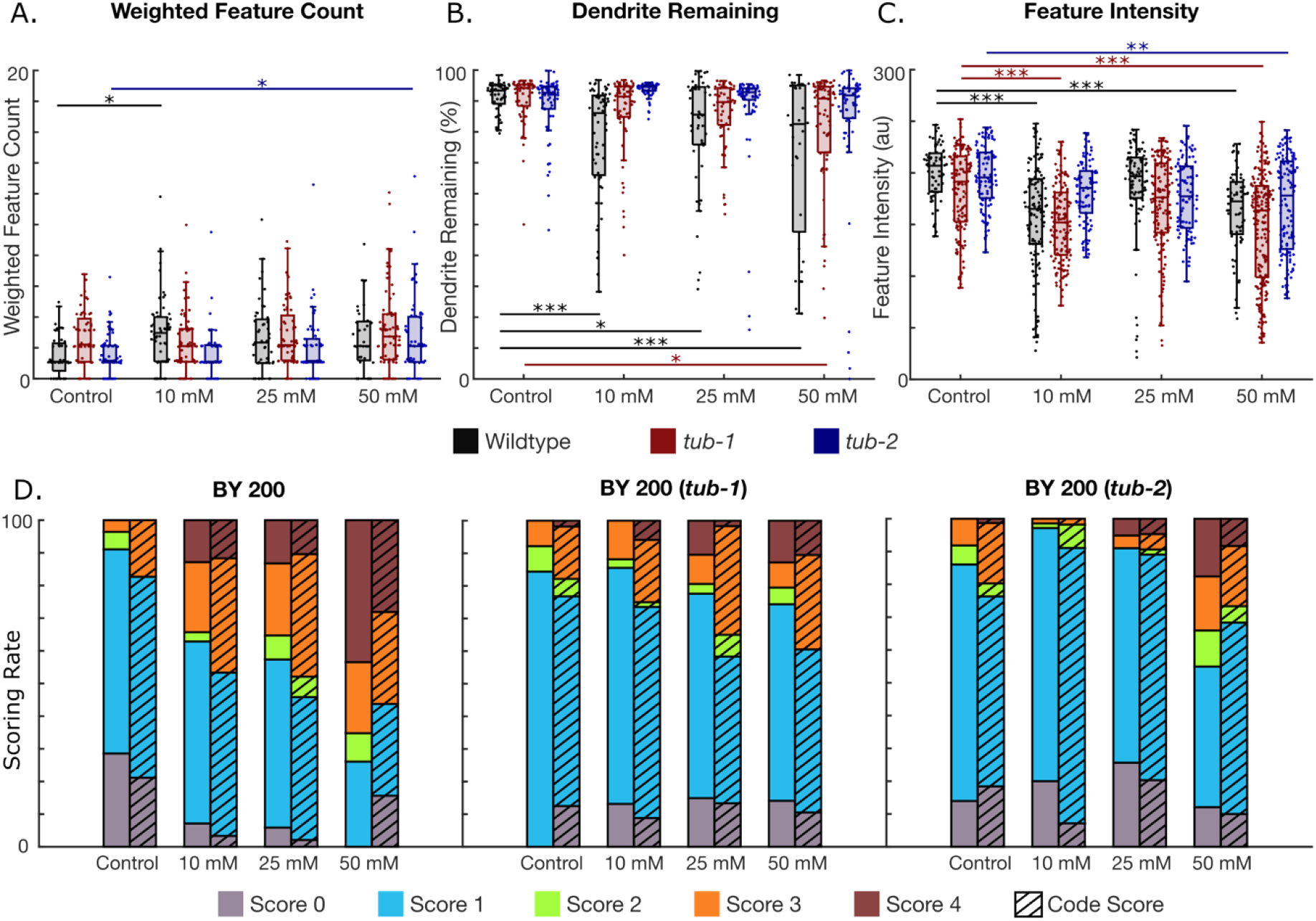
Neurodegeneration quantification of 6-OHDA exposed worms. N=1 represents a single dendrite. A-C. Neurodegeneration metrics for control (N = 52, 56, and 76 for wildtype, tub-1, and tub-2, respectively), 10 mM (N = 60, 68, and 56 for wildtype, tub-1, and tub-2, respectively), 25 mM (N = 48, 56, and 64 for wildtype, tub-1, and tub-2, respectively), and 50 mM (N = 32, 76, and 60 for BY200, tub-1, and tub-2, respectively) 6-OHDA exposures. A. Feature count on dendrites weighted by the amount of dendrite remaining. B. Percentage of dendrite remaining. C. Intensity of detected features. D. Comparison of the neurodegeneration algorithm’s detection compared to manual dendrite scoring. (One-way ANOVA. *p < 0.05; **p < 0.01; ***p < 0.001).

**Fig 6.**
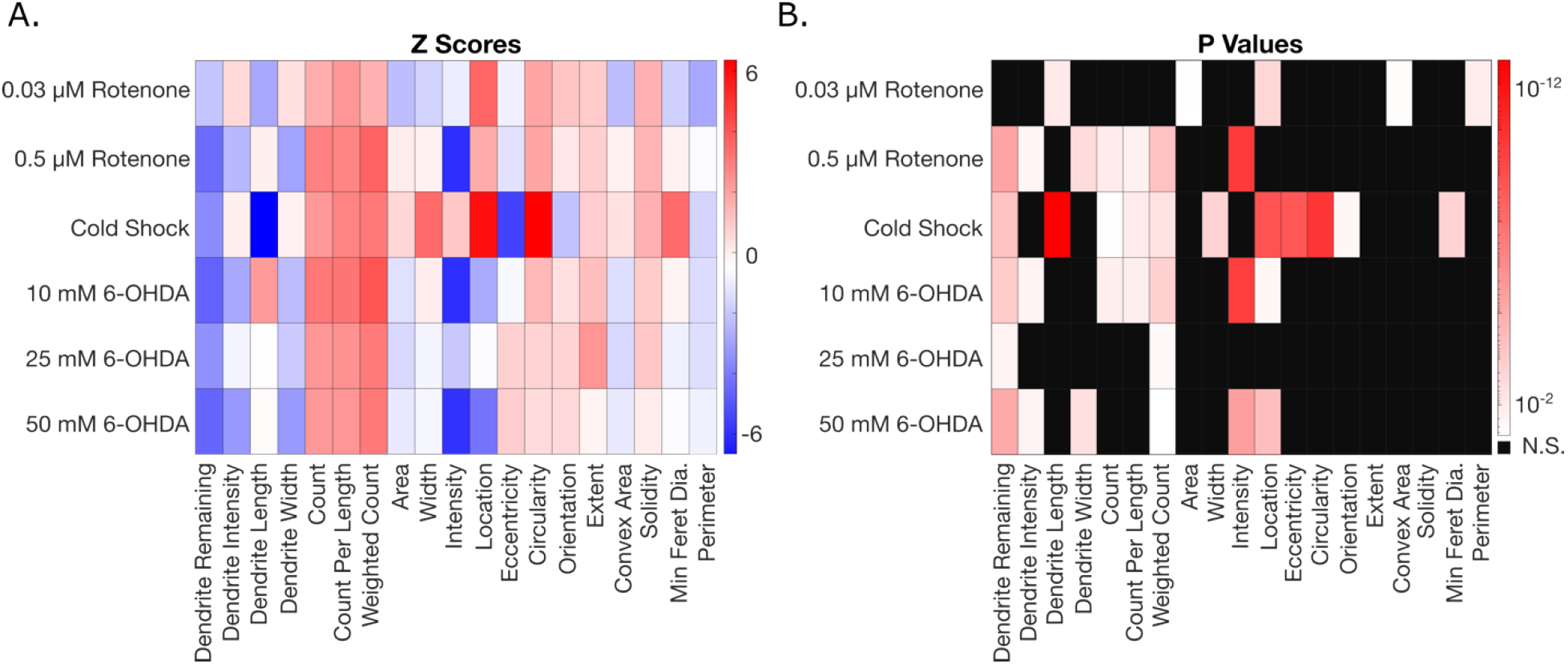
Experimental comparison across different exposure groups compared to their respective control. A. Heat map of z scores for each exposure and its metrics obtained from AUDDIT to show directionality. B. Metrics that were significantly affected for each exposure with p < 0.05. (One-way ANOVA, Dunnett’s multiple comparisons).

### ii. Feature detection

A key morphological feature of dendrite degeneration is the presence of blebbing (sometimes described as “beads on a string”). Blebs are roughly circular, morphological features that protrude from the dendrite(15,36). To identify blebbing features in the dendrite, the algorithm first performs a series of preprocessing steps on the cropped image. First, the image undergoes a local contrast adjustment using the localcontrast() function in MATLAB to improve individual dendrite definition. The extent of contrast enhancement depends on the original cropped image’s background noise and intensity. Next, the image undergoes a 4x resize to increase available pixels for image morphological operations, which helps improve feature detection. The last preprocessing step is a background subtraction to help normalize uneven illumination and reduce noise throughout the image (Fig 2B).

Once the image has been processed, a binarization with a global threshold is utilized to obtain a rough outline of the dendrites. This threshold level is dependent on the noise and overall intensity level of the original image, as image artifacts are more easily detected with noisy or low contrast images. Thus, images with higher noise must have an increased threshold level to minimize the number of unwanted objects placed in the foreground. Since features, or morphological abnormalities on dendrites, are wider than dendrites, horizontal erosion is then performed to remove thin vertical dendrites. The clusters of pixels remaining after the erosion identify the locations of detected morphological features. However, horizontal erosion often causes the shape of resulting pixel clusters to be jagged with a horizontal bias. To improve shape definition of the blebs, an iterative region-growing function, activecontour() using the Chan-Vese method, is utilized at each of these locations (37). Finally, region properties of the bleb features, such as area, circularity, and intensity, are extracted and stored for further analysis (Fig 2B).

### iii. Dendrite tracking

Another important metric in studying dendrite degeneration is dendrite breakage. Tracing the dendrites is challenging as they can break off into different segments, change direction abruptly, and even overlap with one another. Furthermore, while a healthy worm has 4 CEP dendrites, degeneration can result in loss of one or more of them. Our approach begins by starting at the top of the image, and sequentially locating four local maximum intensities and their respective x coordinates at each y value of the image, representing four separate dendrites (Fig 2C). Since dopaminergic neurons may have breaks, some rows in the resulting image will not have all 4 local maxima intensities. Thus, in an attempt to connect each dendrite to itself, the algorithm conducts a morphological closing with a vertical structuring element using the located maximum intensities as foreground objects. These rough dendrite objects are then skeletonized to obtain an improved dendrite track. Next, each tracked object is binned into its respective dendrite index by matching the tracked object’s x location to a running average of dendrite indices. If there is more than one object indexed to the same dendrite at the same y location, the object farther away from the running average will be removed, leaving a maximum of four dendrite indices at each y value. Lastly, each tracked dendrite is smoothed and interpolated to fill in any missing spaces, such that there are four discrete tracks (Fig 2C).

### iv. Degeneration metrics

The last step in the neurodegeneration algorithm includes matching detected features to their respective dendrites, detecting dendrite breakage, and ultimately creating a degeneration metric report for downstream analysis. Bleb features are matched to the corresponding dendrite by comparing track location with the pixel locations of the detected bleb features. Dendrite breakage is detected by sequentially analyzing each pixel row in each individual dendrite track and comparing it to a binarized image of dendrites (S1 Fig). If the tracks do not include any foreground objects from the dendrite binarization, that pixel row of the tracked dendrite would be determined as a break. The final step is compiling the metrics of neurodegeneration into a file for future analysis and visualization. A thorough explanation of all 19 neurodegeneration metrics realized by AUDDIT can be found in S1 Table. We next utilized AUDDIT to score and analyze images of dopaminergic neurons exposed to known sources of dopaminergic degeneration.

### B. Rotenone exposure induces dendritic blebbing, breakage, and neuronal intensity loss

Rotenone is a potent inhibitor of mitochondrial electron transport chain Complex I known to induce oxidative stress(38–40). Despite these dangers, it continues to be used as a pesticide outside of the United States, and as a piscicide inside of the United States(40). Over two decades of research has shown that rotenone can consistently induce dopaminergic degradation both in cell cultures and live organisms(41–45). We aimed to use our algorithmic platform to gain further quantitative insight into neuronal deterioration induced by rotenone exposure in *C. elegans*. Examples of healthy, moderately degenerated, and severe degeneration induced by control, 0.03 µM and 0.5 µM rotenone exposures can be seen in Fig 3A. The rectangles overlaid on top of the images depict the location of features detected by the algorithm.

We extracted various metrics of neurodegeneration, such as feature count (blebbing), dendrite length, dendrite remaining, and dendrite intensity. The extracted data from each dendrite labeled with a star, rectangle, triangle, or circle in the example images in Fig 5A is shown in Fig 3B-D. Worms exposed to 0.5 µM rotenone exhibited significantly higher rates of blebbing per dendrite length than control worms (Fig 3B). Interestingly, AUDDIT revealed that worms exposed to 0.03 µM rotenone have shorter dendrites than controls (Fig 3C). We hypothesize that this is due to rotenone slowing development in worms, and thus making the dendrites shorter(46). The worms exposed to 0.5 µM rotenone were grown for an extra day to account for slower development, which permitted the dendrites enough time to reach the same length as the controls. In addition, worms exposed to 0.5 µM rotenone also showed an increase in dendrite breakage (Fig 3D). Interestingly, the 0.5 µM dosed worms exhibited significantly dimmer dendrites than the controls (Fig 3E). This may suggest that a decrease in fluorescent intensity driven by a *dat-1* promoter can be used as a marker of neurodegeneration. However, decreased dendrite intensity may also reflect decreased GFP protein production due to decreased energetics. The remaining neurodegeneration metrics for rotenone exposed worms can be found in S2 Fig. Ultimately, the neurodegeneration algorithm’s neurodegeneration report is consistent with previous literature showing that rotenone induces neurodegeneration in *C. elegans*(15).

### C. Cold Shock induces large, dispersed dendritic blebs

Acute cold shock has been shown to be harmful to worms, with various negative phenotypes being reported(47,48). Previous experiments have shown the prevalence of dendritic blebbing in the PVD neurons of worms following cold shock(22). For this reason, we were interested in utilizing our algorithm to see if cold shock induced blebbing features in the CEP neurons as well. Thus, we exposed worms with fluorescently tagged CEP neurons to a 16-hour cold shock at 4 °C. Example images of pre-cold shock, control, and cold shocked worms, as well as the location of features detected by the algorithm, are shown in Fig 4A. Interestingly, cold shocking worms for 16 hours significantly increases the number of features (blebs) on the dendrites (Fig 4B). Moreover, detected features of cold shocked worms were located significantly closer to the cell body and more dispersed throughout the dendrite (Fig 4C). Cold shocked worms also exhibited larger features (Fig 4D). Curiously, 16 hour aged controls and cold shocked worms exhibited features with smaller extents compared to younger, pre-cold shocked worms (Fig 4E). Extent is another metric to describe a feature’s shape, with bounds of 0 and 1. An extent of 1 refers to a filled rectangle, while an extent of approximately 0 refers to a cross. Thus, the algorithm detected that older control and cold shocked worms have features with more concave and variable shapes. In addition to features, cold shock induced a significant increase in dendritic breakage compared to both pre-cold shock and control worms (Fig 4F). The other metrics extracted by AUDDIT for the cold shock experiments can be found in S3 Fig. Ultimately, the neurodegeneration algorithm revealed that cold shock induces dopaminergic degeneration, mainly by increasing the number and size of bleb-like features in *C. elegans*.

### D. 6-OHDA-induced dendritic breakage is minimized in *tub-1, tub-2* mutants

To test the algorithm’s versatility, we tested dopaminergic neurodegeneration after challenge with 6-OHDA in different genetic backgrounds, and with a different imaging set up. 6-OHDA is known to induce oxidative stress via free radicals and inhibition of mitochondrial electron transport chain Complex I and Complex IV(49). Thus, we exposed wildtype nematodes (BY200) with no mutations, as well worms with deletions in *tub-1* and *tub-2*, to 10 mM, 25 mM, and 50 mM 6-OHDA to induce dopaminergic degeneration. *Tub-1* and t*ub-2* are homologs of human *TUBBY* proteins that have previously been shown to increase fat storage. However, in our quantification, *tub-1* mutants showed decreased fat storage, whereas *tub-2* mutants showed increased fat storage (S4 Fig). Images of the worms’ neurons were taken at a lower magnification and with a different camera than the images acquired for the rotenone and cold shock experiments. Remarkably, AUDDIT was able to extract the same quantitative metrics by only changing the input pixel size. Interestingly, the weighted feature count, or the features per dendrite remaining, did not change between most groups. There was a significant increase in blebbing features between wildtype unexposed worms and the 10 mM dosed worms, as well as the *tub-2* mutant controls and the 50 mM dosed worms (Fig 5A). Moreover, dendrite breakage seems to be a significant marker of neurodegeneration in wildtype worms when exposed to 6-OHDA. Interestingly, the *tub-1* and *tub-2* mutations appear to be protective to dendrite breakage caused by 6-OHDA exposure, with the *tub-2* mutation possibly providing more protection (Fig 5B). This indicates that the quantity of lipids present in the worm does not alter susceptibility to neurodegeneration, but that tubby-like proteins or related signaling pathways may play a role in 6-OHDA induced neurodegeneration. Another interesting finding, which would not be possible to detect through manual neurodegeneration scoring, is that feature pixel intensity decreases with higher 6-OHDA exposures in all worm strains (Fig 5D). Again, the *tub-*2 mutation appears to be protective against feature intensity loss, as only the highest exposure concentration resulted in significant differences. The remaining neurodegeneration metrics detected by the algorithm can be seen in S5 Fig.

To verify the robustness of the neurodegeneration algorithm, we sought to compare results of the code to the current state-of-the-art manual scoring method with images of 6-OHDA exposure (Fig 5D). The images were manually scored as follows: no degeneration (0), blebs on less than half of the dendrite (1), blebs on more than half of the dendrite (2), less than 50% breakage (3), and more than 50% breakage (4). We then binned the results from the code using the following procedure: more than 85% dendrite remaining and no blebs (0), less than 5 blebs (1), more than 5 blebs (2), between 50% and 85% dendrite remaining (3), and less than 50% dendrite remaining (4). The neurodegeneration algorithm is consistent with current manual methods for scoring dopaminergic neurodegeneration, which shows its readiness to be used in future dopaminergic neurodegeneration studies.

### E. AUDDIT enables comparative degenerative analysis between exposures

After individually quantifying degeneration of rotenone, cold shock, and 6-OHDA separately, we sought to investigate whether different exposures result in distinct degeneration patterns. We quantified the z-score for neurodegeneration metrics of each treatment compared to their control. These z-scores along with its associated p-value are shown as heatmaps in Fig 6. It becomes clear each stressor causes similar directional effects for most metrics besides intensity and length, yet significance values differ between exposures. Interestingly, cold shock appears to affect feature morphology more than rotenone and 6-OHDA, as a cold shock exposure makes features larger and more circular when compared to their control. Alternatively, this difference could be a result of the improved resolution and reduced noise from confocal images as compared to those obtained with 63x epifluorescence (rotenone experiments) and 40x epifluorescence (6-OHDA experiments).

Most exposures significantly decreased the amount of dendrite remaining, which indicates that dendrite breakage is a key metric in dopaminergic neuron degeneration. Dendrite and feature intensities, which would not be measurable through manual scoring, may also be important when attempting to differentiate degeneration. Interestingly, both rotenone and 6-OHDA significantly decrease both intensity measurements, while the cold shock exposure has no effect. This indicates that the chemical exposures to rotenone and 6-OHDA had a greater effect on the overall fluorescence of the neuron rather than feature morphology.

Moreover, both 0.5 µM rotenone and 10 mM 6-OHDA have similar degenerative effects in both significance and directionality. It is important to note that since each experiment was performed on worms of varying ages and culture conditions, definitive conclusions must be taken with caution. Thus, a more in-depth comparison of degeneration patterns resulting from exposures should be completed by standardizing worm age, culture conditions, and imaging setups. More broadly, however, these results indicate that the metrics obtained from AUDDIT have the potential to elucidate differences in degeneration patterns between different exposures, particularly if other parameters (eg, microscopy) are held constant.

## IV. Discussion

We have built an algorithmic platform that is able to analyze large sets of images from *C. elegans* CEP dopaminergic neurons to produce quantitative information. The exclusion of machine learning permits it to be used on images taken with different microimaging setups without additional training. To test its versatility and robustness, we analyzed images of fluorescently tagged CEP from multiple imaging setups, including different cameras, microscopes, and capturing resolutions. While this platform was built using MATLAB, we have created a standalone app with an intuitive user interface that requires no technical knowledge to use. The app asks the user to specify the imaging settings (camera pixel size), then analyzes batches of images and sends the results to a user specified directory. The app, as well as the MATLAB functions and scripts, can be found in the supplementary information and are available at: https://github.com/zkalmanson/AUDDIT. The results shown in this publication used AUDDIT v1.0. We plan to continue to improve AUDDIT with performance updates as needed and will add the new versions to the GitHub repository.

Using this algorithm, we were able to demonstrate and investigate three different biological phenomena. First, our evaluation of the established neurotoxicant and Complex I inhibitor rotenone was cohesive with previous literature in showing an increase in blebbing and dendrite breakage in *C. elegan*s. However, it expands on this body of work by further uncovering that even 0.03 µM exposure results in shorter dendrites than controls, and 0.5 µM rotenone exposure causes a decrease in the brightness of dendrites. Decreased dendrite intensity could reflect a neurodegenerative feature or even decreased GFP protein production due to decreased energetics. Second, our investigation of the impact of cold shock demonstrates that beyond detection of blebs, additional characteristics about the dendrites can be observed. For example, we found that dendrites from cold shocked worms not only had more blebs, but they exhibited wider shaped blebs than their control counterparts. Providing a sharp contrast to rotenone exposure, the significant increase in blebbing due to cold shock suggests a distinct damage pattern than the breakage induced by rotenone exposure. This difference is one that would not be as easily detected by manual scoring and may not be detected at all by brightness quantification. Finally, we utilized another canonical dopaminergic neurotoxicant, 6-OHDA, and two strains with mutations in tubby-like proteins to simultaneously evaluate the impact of two genetic mutations and their alteration to 6-OHDA susceptibility. We show that although one mutation increases fat storage and the other decreases it, both are protective from 6-OHDA exposure. Neither mutation alone results in increased neurodegeneration. Thus, deletions in tubby-like proteins, regardless of the subsequent impact on lipid burden within the worms, were protective. Modifying the discussion of obesity and neurodegenerative disorders, this suggests increased adiposity alone did not increase susceptibility to neurodegeneration(50). Together, these three investigations confirm the ability of AUDDIT to quantify known impacts of neurotoxicants, identify induction of neurodegeneration by previously unknown sources, and work in conjunction with genetic mutations to allow for more complex analyses.

One limitation of AUDDIT is that there can only be one worm in the frame of the image for the cropping feature to work accurately. This should be considered by the user when imaging worms. Moreover, the algorithm requires all four CEP dendrites to be visible. If dendrites overlap, the algorithm will detect the overlapping region as breaks, which may skew results. Good and bad example input images for AUDDIT can be visualized in S6 Fig. Since the tracking algorithm starts from one end of the image and moves forward, this platform struggles to track dendrites that exhibit kinks that would require reversing direction (greater or equal than 90 degrees). If a dendrite makes a turn greater than 90 degrees and momentarily moves backwards, the tracking algorithm will not follow it, but rather interpolate to the next point forward towards the distal end of the dendrite. The smoothing features of the algorithm help to prevent these erratic jumps from confusing the tracking, but in the process will end up smoothing out the kink of the dendrite if the kink is greater than or equal to 90 degrees. We plan to address some of these limitations in future versions of AUDDIT which will be continually added to the GitHub project.

An advantage of this algorithmic approach is the ability to obtain quantitative metrics for subtle markers of dopaminergic degeneration from many images without user intervention.

Manual scoring can be very time-consuming and is also susceptible to inconsistent scoring even with blinding. For example, researchers may subconsciously score a moderately damaged dendrite as more degenerated if they just scored many healthy dendrites in a row. In these instances, particularly important for large data sets, a uniform evaluation system is critical. Moreover, a major advantage of AUDDIT lies in its ability to capture quantitative information about the dendrite. For instance, the high dose rotenone worms showing lower intensity features was a metric that would not have been able to be scored by the human eye. Likewise, while 6-OHDA has been shown to induce dopaminergic deterioration in worms(13,14,16), the studies have been limited by discrete, manual scoring systems. While this method of scoring is adequate for obtaining data like the percent of population experiencing deterioration, it provides less information about the exact degree of damage shown by each individual worm. The algorithm can detect 19 different metrics of neurodegeneration, which can provide insight into how different exposures affect dopaminergic neurons. For example, rotenone exposure and cold shock produce significant increases in feature counts (blebs), while 6-OHDA mainly induces dendrite breakage. Finally, because this analysis approach generates continuous rather than categorical data, the data is more amenable to parametric statistical analysis than standard common categorical score-based approaches.

Considering the progressive nature of many neurodegenerative diseases, quantitative information using multiple metrics would also be a powerful tool for observing changes in neuronal health over time. This could help provide key insights into how toxicants or genetic defects may exacerbate neuronal loss in an aging organism. The *C. elegans* nematode shows great promise as a model for studying dopaminergic degradation. We have constructed AUDDIT to help standardize the analysis of scoring dopaminergic images and minimize bias. Further recommendations would be to use this platform as a tool for comparing the degradative effects and patterns for a variety of different toxins under standardized culturing and imaging conditions.

Looking at multiple quantitative metrics side by side can provide a more comprehensive understanding of the mechanisms underlying these chemicals. In addition, observing changes in dopaminergic deterioration over a worm’s lifespan could advance our current model by providing insight into the progression of the degradative effects, rather than just snapshots. Ultimately, AUDDIT provides an unbiased and faster alternative to current, manual dopaminergic degeneration scoring in *C. elegans*, paving the way for higher throughput studies.

## V. Acknowledgements

We would like to acknowledge and thank the *C. elegans* Gene Knockout Project at the Oklahoma Research Foundation, a part of the International *C. elegans* Gene Knockout Consortium, for providing the strains RB1600, *tub-1*(ok192), and RB2237, *tub-2*(ok3024). This work was funded in part by NSF awards 1838314 and 1947498 (ASM) and NIH award K99-ES029552 (JHH).

## References

1. Poewe W, Seppi K, Tanner CM, Halliday GM, Brundin P, Volkmann J, et al. Parkinson disease. Nat Rev Dis Primers. 2017 Mar 23;3:1–21.

2. Tatton WG, Chalmers-Redman R, Brown D, Tatton N, Schapira, Hunot, et al. Apoptosis in Parkinson’s disease: Signals for neuronal degradation. vol. 53, Annals of Neurology. 2003.

3. Vázquez-Vélez GE, Zoghbi HY, Duncan D. Annual Review of Neuroscience Parkinson’s Disease Genetics and Pathophysiology. Annu Rev Neurosci [Internet]. 2021;44:87–108. Available from: https://doi.org/10.1146/annurev-neuro-100720-

4. Rocha EM, Keeney MT, di Maio R, de Miranda BR, Greenamyre JT. LRRK2 and idiopathic Parkinson’s disease. vol. 45, Trends in Neurosciences. Elsevier Ltd; 2022. p. 224–36.

5. Tanner CM, Kame F, Ross GW, Hoppin JA, Goldman SM, Korell M, et al. Rotenone, paraquat, and Parkinson’s disease. Environ Health Perspect. 2011 Jun;119(6):866–72.

6. Bernhardt ES, Rosi EJ, Gessner MO. Synthetic chemicals as agents of global change. Front Ecol Environ. 2017 Mar 1;15(2):84–90.

7. Judson R, Richard A, Dix DJ, Houck K, Martin M, Kavlock R, et al. The toxicity data landscape for environmental chemicals. vol. 117, Environmental Health Perspectives. 2009. p. 685–95.

8. Falkenburger BH, Schulz JB. Limitations of cellular models in Parkinson’s disease research. J Neural Transm. 2006;70:261–8.

9. Nass R, Hall DH, Miller III DM, Blakely RD. Neurotoxin-induced degeneration of dopamine neurons in Caenorhabditis elegans. PNAS [Internet]. 2002;99(5):3264–9. Available from: www.pnas.org

10. Cao P, Yuan Y, Pehek EA, Moise AR, Huang Y, Palczewski K, et al. Alpha-synuclein disrupted dopamine homeostasis leads to dopaminergic neuron degeneration in Caenorhabditis elegans. PLoS One. 2010 Feb 19;5(2).

11. Dawson TM, Ko HS, Dawson VL. Genetic Animal Models of Parkinson’s Disease. vol. 66, Neuron. 2010. p. 646–61.

12. Benedetto A, Au C, Avila DS, Milatovic D, Aschner M. Extracellular dopamine potentiates Mn-induced oxidative stress, lifespan reduction, and dopaminergic neurodegeneration in a BLI-3-dependent manner in caenorhabditis elegans. PLoS Genet. 2010 Aug;6(8).

13. Offenburger SL, Gartner A. 6-hydroxydopamine (6-OHDA) Oxidative Stress Assay for Observing Dopaminergic Neuron Loss in Caenorhabditis elegans. Bio Protoc. 2018;8(18).

14. Hartman JH, Gonzalez-Hunt C, Hall SM, Ryde IT, Caldwell KA, Caldwell GA, et al. Genetic defects in mitochondrial dynamics in caenorhabditis elegans impact ultraviolet c radiation- and 6-hydroxydopamine-induced neurodegeneration. Int J Mol Sci. 2019 Jul 1;20(13).

15. Bijwadia SR, Morton K, Meyer JN. Quantifying Levels of Dopaminergic Neuron Morphological Alteration and Degeneration in Caenorhabditis elegans. Journal of Visualized Experiments. 2021 Nov 1;2021(177).

16. Offenburger SL, Ho XY, Tachie-Menson T, Coakley S, Hilliard MA, Gartner A. 6-OHDA-induced dopaminergic neurodegeneration in Caenorhabditis elegans is promoted by the engulfment pathway and inhibited by the transthyretin-related protein TTR-33. PLoS Genet. 2017;14(1).

17. Mor DE, Sohrabi S, Kaletsky R, Keyes W, Tartici A, Kalia V, et al. Metformin rescues Parkinson’s disease phenotypes caused by hyperactive mitochondria. PNAS [Internet]. 2020;117(42):26438–16447. Available from: www.pnas.org/cgi/doi/10.1073/pnas.2009838117

18. Jadiya P, Khan A, Sammi SR, Kaur S, Mir SS, Nazir A. Anti-Parkinsonian effects of Bacopa monnieri: Insights from transgenic and pharmacological Caenorhabditis elegans models of Parkinson’s disease. Biochem Biophys Res Commun. 2011 Oct 7;413(4):605–10.

19. Yuan Y, Cao P, Smith MA, Kramp K, Huang Y, Hisamoto N, et al. Dysregulated LRRK2 signaling in response to endoplasmic reticulum stress leads to dopaminergic neuron degeneration in C. elegans. PLoS One. 2011;6(8).

20. Smith LL, Ryde IT, Hartman JH, Romersi RF, Markovich Z, Meyer JN. Strengths and limitations of morphological and behavioral analyses in detecting dopaminergic deficiency in Caenorhabditis elegans. Neurotoxicology. 2019 Sep 1;74:209–20.

21. Gaeta AL, Brucker Nourse J, Willicott K, McKay LE, Keogh CM, Peter K, et al. Systemic RNA Interference Defective (SID) genes modulate dopaminergic neurodegeneration in C. elegans. PLoS Genet. 2022 Aug 19;18(8).

22. Saberi-Bosari S, Flores KB, San-Miguel A. Deep learning-enabled analysis reveals distinct neuronal phenotypes induced by aging and cold-shock. BMC Biol. 2020 Sep 23;18(1).

23. Cothren S, Meyer J, Hartman J. Blinded Visual Scoring of Images Using the Freely-available Software Blinder. Bio Protoc. 2018;8(23).

24. Chothani P, Mehta V, Stepanyants A. Automated tracing of neurites from light microscopy stacks of images. Neuroinformatics. 2011 Sep;9(2–3):263–78.

25. Liu S, Zhang D, Song Y, Peng H, Cai W. Automated 3-D Neuron Tracing with Precise Branch Erasing and Confidence Controlled Back Tracking. IEEE Trans Med Imaging. 2018 Nov 1;37(11):2441–52.

26. Zhou Z, Kuo HC, Peng H, Long F. DeepNeuron: an open deep learning toolbox for neuron tracing. Brain Inform. 2018 Dec 1;5(2).

27. Huang Q, Cao T, Chen Y, Li A, Zeng S, Quan T. Automated Neuron Tracing Using Content-Aware Adaptive Voxel Scooping on CNN Predicted Probability Map. Front Neuroanat. 2021 Aug 23;15.

28. Boyd WA, Smith M v., Freedman JH. Caenorhabditis elegans as a model in developmental toxicology. Methods in Molecular Biology. 2012;889:15–24.

29. Jobson MA, Jordan JM, Sandrof MA, Hibshman JD, Lennox AL, Baugh LR. Transgenerational effects of early life starvation on growth, reproduction, and stress resistance in Caenorhabditis elegans. Genetics. 2015 Sep 1;201(1):201–12.

30. Stiernagle T. Maintenance of C. elegans. WormBook : the online review of C. elegans biology. 2006. p. 1–11.

31. Chuang‐Stein C, Tong DM. Multiple comparisons procedures for comparing several treatments with a control based on binary data. Stat Med. 1995;14(23):2509–22.

32. Dunnett CW. A Multiple Comparison Procedure for Comparing Several Treatments with a Control. vol. 50, Source: Journal of the American Statistical Association. 1955.

33. Albrecht PA, Fernandez-Hubeid LE, Deza-Ponzio R, Martins AC, Aschner M, Virgolini MB. Developmental lead exposure affects dopaminergic neuron morphology and modifies basal slowing response in Caenorhabditis elegans: Effects of ethanol. Neurotoxicology. 2022 Jul 1;91:349–59.

34. Ke T, Santamaria A, Rocha JBT, Tinkov A, Bornhorst J, Bowman AB, et al. Cephalic Neuronal Vesicle Formation is Developmentally Dependent and Modified by Methylmercury and sti-1 in Caenorhabditis elegans. Neurochem Res. 2020 Dec 1;45(12):2939–48.

35. Vozdek R, Pramstaller PP, Hicks AA. Functional Screening of Parkinson’s Disease Susceptibility Genes to Identify Novel Modulators of α-Synuclein Neurotoxicity in Caenorhabditis elegans. Front Aging Neurosci. 2022 Apr 27;14.

36. Chen S, Tran S, Sigler A, Murphy TH. Automated and quantitative image analysis of ischemic dendritic blebbing using in vivo 2-photon microscopy data. J Neurosci Methods. 2011 Feb 15;195(2):222–31.

37. Chan TF, Vese LA. Active Contours Without Edges. vol. 10, IEEE TRANSACTIONS ON IMAGE PROCESSING. 2001.

38. Li N, Ragheb K, Lawler G, Sturgis J, Rajwa B, Melendez JA, et al. Mitochondrial complex I inhibitor rotenone induces apoptosis through enhancing mitochondrial reactive oxygen species production. Journal of Biological Chemistry. 2003 Mar 7;278(10):8516–25.

39. Heinz S, Freyberger A, Lawrenz B, Schladt L, Schmuck G, Ellinger-Ziegelbauer H. Mechanistic Investigations of the Mitochondrial Complex i Inhibitor Rotenone in the Context of Pharmacological and Safety Evaluation. Sci Rep. 2017 Apr 4;7.

40. Lawana V, Cannon, JR. Chapter Five - Rotenone neurotoxicity: Relevance to Parkinson’s disease. In: Aschner M, Costa, LG, editor. Neurotoxicity of Pesticides. Elsevier; 2020. pp. 209–254.

41. Testa CM, Sherer TB, Greenamyre JT. Rotenone induces oxidative stress and dopaminergic neuron damage in organotypic substantia nigra cultures. Molecular Brain Research. 2005 Mar 24;134(1):109–18.

42. Alam M, Schmidt WJ. Rotenone destroys dopaminergic neurons and induces parkinsonian symptoms in rats. Behavioral Brain Research [Internet]. 2002;317–24. Available from: www.elsevier.com/locate/bbr

43. Coulom H, Birman S. Chronic exposure to rotenone models sporadic Parkinson’s disease in Drosophila melanogaster. Journal of Neuroscience. 2004 Dec 1;24(48):10993–8.

44. Sasajima H, Miyazono S, Noguchi T, Kashiwayanagi M. Intranasal Administration of Rotenone to Mice Induces Dopaminergic Neurite Degeneration of Dopaminergic Neurons in the Substantia Nigra. vol. 108, Biol. Pharm. Bull. 2017.

45. Norazit A, Meedeniya ACB, Nguyen MN, MacKay-Sim A. Progressive loss of dopaminergic neurons induced by unilateral rotenone infusion into the medial forebrain bundle. Brain Res. 2010 Nov 11;1360:119–29.

46. Mello DF, Bergemann CM, Fisher K, Chitrakar R, Bijwadia SR, Wang Y, et al. Rotenone Modulates Caenorhabditis elegans Immunometabolism and Pathogen Susceptibility. Front Immunol. 2022 Feb 22;13.

47. Robinson JD, Powell JR. Long-term recovery from acute cold shock in Caenorhabditis elegans. BMC Cell Biol. 2016 Jan 12;17(1):1–11.

48. Jiang W, Wei Y, Long Y, Owen A, Wang B, Wu X, et al. A genetic program mediates cold-warming response and promotes stress-induced phenoptosis in C. elegans. Elife [Internet]. 2018; Available from: https://doi.org/10.7554/eLife.35037.001

49. Glinka Y, Gassen M, Youdim MBH. Mechanism of 6-hydroxydopamine neurotoxicity. Journal of Neural Transmission . 1997;(50):55–66.

50. Mazon JN, de Mello AH, Ferreira GK, Rezin GT. The impact of obesity on neurodegenerative diseases. vol. 182, Life Sciences. Elsevier Inc.; 2017. p. 22–8.

